# Bacteria induce an amoeboid phase in coccolithophores that persists after bloom collapse

**DOI:** 10.1101/2025.01.06.631481

**Authors:** Sophie T. Zweifel, Richard J. Henshaw, Oliver Müller, Johannes M. Keegstra, Roberto Pioli, Clara Martínez-Pérez, Uria Alcolombri, Estelle Clerc, Roman Stocker

**Author notes:** For correspondence/requests for materials. **Author Contributions:** S.T.Z., R.J.H., and R.S. designed the study; S.T.Z. performed and analyzed amoeboid induction experiments, bacterial characterization, light and electron microscopy imaging, EDX analysis, and flow cytometry experiments; O.M. carried out PAM measurements and analyzed the resulting data; S.T.Z. and R.P. conducted fluorescence and time-lapse microscopy on amoeboid cells; S.T.Z. and J.M.K. carried out single-cell tracking analysis; R.J.H. performed tracking analysis of the amoeboid ellipticity transformation; S.T.Z., R.J.H., E.C and R.S. discussed the results; and S.T.Z., R.J.H. and R.S. wrote the manuscript with contributions from all other authors. **Competing Interest Statement:** The authors declare no competing interests.

## Abstract

Coccolithophores, who contribute approximately 1-10% of phytoplankton biomass, are crucial players in the ocean’s biogeochemical cycles. Significant contributions come from bloom-forming species including *Gephyrocapsa huxleyi* (formerly *Emiliania huxleyi),* which has served as a model system for investigating algal-bacterial and agal-viral interactions as well as algal fitness responses to environmental changes. Coccolithophores follow a biphasic lifecycle, existing as motile haploid and non-motile diploid phases. Here we characterize a third, ‘amoeboid’ phase, using a combination of light, electron and phase amplitude modulation (PAM) microscopy. Using time-resolved imaging we captured the rapid morphological transition from a spherical haploid to an elongated motile amoeboid cell. Cell tracking revealed slower and more directional swimming compared to haploid cells. Amoeboid metamorphosis was triggered by exposure to bacteria, including strains isolated from *G. huxleyi* blooms and known *G. huxleyi* pathogens, but not by a range of classical phytoplankton stressors, including viral infection and oxidative stress. Further, the sub-population of haploids which switched to the amoeboid phase persisted past the rapid crash of the haploid population. This amoeboid phase was only observed in the primary bloom-forming coccolithophore species *G. huxleyi* and *Geopharycapsa oceanica* in stationary phase when exposed to high bacterial concentrations, typical of late-stage algal bloom events. Photophysiology of amoeboid cells was confirmed to be unaltered via PAM microscopy, indicating these cells are metabolically active. These findings highlight a previously uncharacterized morphotype in this important phytoplankton species and suggest that the amoeboid phase could be a bacteria-resistant morphotype following algal bloom collapse.

## Introduction

Phytoplankton are responsible for about 50% of global primary production^1,2^ and are thus a crucial component of ecosystem-scale biogeochemical processes. Prominent amongst the phytoplankton community are coccolithophores, a group of calcifying microalgae that account for nearly 10% of phytoplankton biomass^3,4^ and are characterized by an intricate calcium carbonate shell comprised of overlapping coccoliths. These autotrophs are vital components of both the organic and inorganic carbon pumps, sequestering over a billion tons of CaCO_3_ annually in the deep ocean upon cell death^5,6^ and composing key links in biogeochemical cycling and ocean alkalinity dynamics^7^. Given their global prominence, coccolithophores have been established as a model system to study a diverse range of marine processes, from physiological responses to ocean acidification in modern and historic oceans^8–10^ to microscale processes including algal interactions with pathogenic bacteria and viruses, which have been linked to the termination of coccolithophore blooms through release of algicidal compounds and by triggering programmed cell death^11–14^.

In contrast to some other phytoplankton such as diatoms^15^, coccolithophores follow a biphasic lifecycle alternating between a diploid morphotype that is non-motile and calcified, and a vegetatively growing haploid morphotype that is bi-flagellated, motile and minimally calcified^16–21^. Two coccolithophores, *Geopharycapsa huxleyi* (formerly known as *Emiliania huxleyi*^22^) and *Geopharycapsa oceanica,* ubiquitous in the euphotic zone in almost all ocean regions^23,24^, often form kilometer-scale blooms that are visible from space^23,25–27^. These two species alone account for more than a third of oceanic calcium carbonate formation^28^. A critical component to shortening *G. huxleyi* blooms or causing their collapse is a rise in large virus-like particles (LVLPs), now known as *G. huxleyi* viruses (EhVs)^29–32^, which can trigger shifts in metamorphosis and ploidy of a subpopulation of the virally susceptible diploid into a virally resistant flagellated haploid or diploid morphotype^33–35^. In the case of *G. huxleyi*, a third life phase has been anecdotally observed^36,16,17,37^. This ‘amoeboid’ morphotype was first reported as a fusiform-shaped cell with ‘wriggling’ motility^36^. A later study found these amoeboid cells within stationary cultures and proposed they had lost their cell wall^16^. Beyond these reports, this elusive morphotype has remained unexplored, in particular as to its triggers and ecological function.

Here we report that certain *G. huxleyi*-associated bacteria trigger rapid metamorphosis in haploid *G. huxleyi* cells to this amoeboid morphotype. After bacterial exposure, cells from late-stage cultures transform within minutes into a motile, non-calcified, amoeboid with apparent bacterial resistance. We extensively characterize this amoeboid phase, which has only anecdotally been reported before^16,36–38^, using a combination of flow cytometry and time-resolved video-, electron and phase modulation microscopy. We identified the environmental conditions responsible for triggering the metamorphosis into the amoeboid phase, finding them to be compatible with those characteristic of the demise of algal blooms. Strikingly, in contrast to haploids, amoeboids are observed to persist throughout exposure to harmful bacterial conditions. Our results indicate that the amoeboid metamorphosis could be a survival response: transforming into a bacteria-resistant morphotype might enable cells to persist beyond the collapse of a bloom, a further step in a shape transition cascade starting from the diploid to haploid transformation under viral exposure^13,14,29,32,35^. The amoeboid phase may thus represent an important life cycle stage in the bloom dynamics of coccolithophores, furthering our understanding of the complex processes underpinning algal blooms.

## Results

### Haploid bloom-forming coccolithophores can undergo a rapid morphological transition to a motile ‘amoeboid’ phase

Using time-lapse microscopy, we observed a previously uncharacterized morphotype amongst xenic haploid cultures of the model coccolithophore *G. huxleyi* (RCC1217), whereby the cell rapidly elongates from the typical spherical shape to an ellipsoidal form that we hereafter refer to as the amoeboid phase (Fig. 1A, Movie S1). This phase has previously been briefly reported ^16,36–38^, where elongated cells were observed to undergo a “wriggling” motion^36^. Electron microscopy imaging later captured an image of an elongated cell body^16^, yet neither a detailed characterization nor the origin of this phase were pursued. Using single-cell tracking and image analysis (Fig. 1B), we found that the transition occurs rapidly, with the cell’s eccentricity dramatically increasing over a 15 min period before stabilizing. The cell ceases motility during the transformation and resumes it thereafter (Movie S2). Scanning electron microscopy (SEM, Methods) images of the haploid (Fig. 1C) and amoeboid (Fig. 1D) cells confirm the stark contrast in morphology between the two phases. Using energy-dispersive X-ray spectroscopy (EDX, Methods), we discovered that the amoeboid phase lacks calcification, evidenced by the absence of characteristic calcium carbonate EDX peaks (SI Fig. S1). Microbial adhesion to hydrocarbons assay revealed a 39 ± 2% increase in surface hydrophobicity in the amoeboid phase compared to the haploid phase (Methods; SI Fig. S2). Such cell surface modifications can alter cell recognition as well as enhance stickiness and cell aggregation^39,40^.

**Fig. 1.**
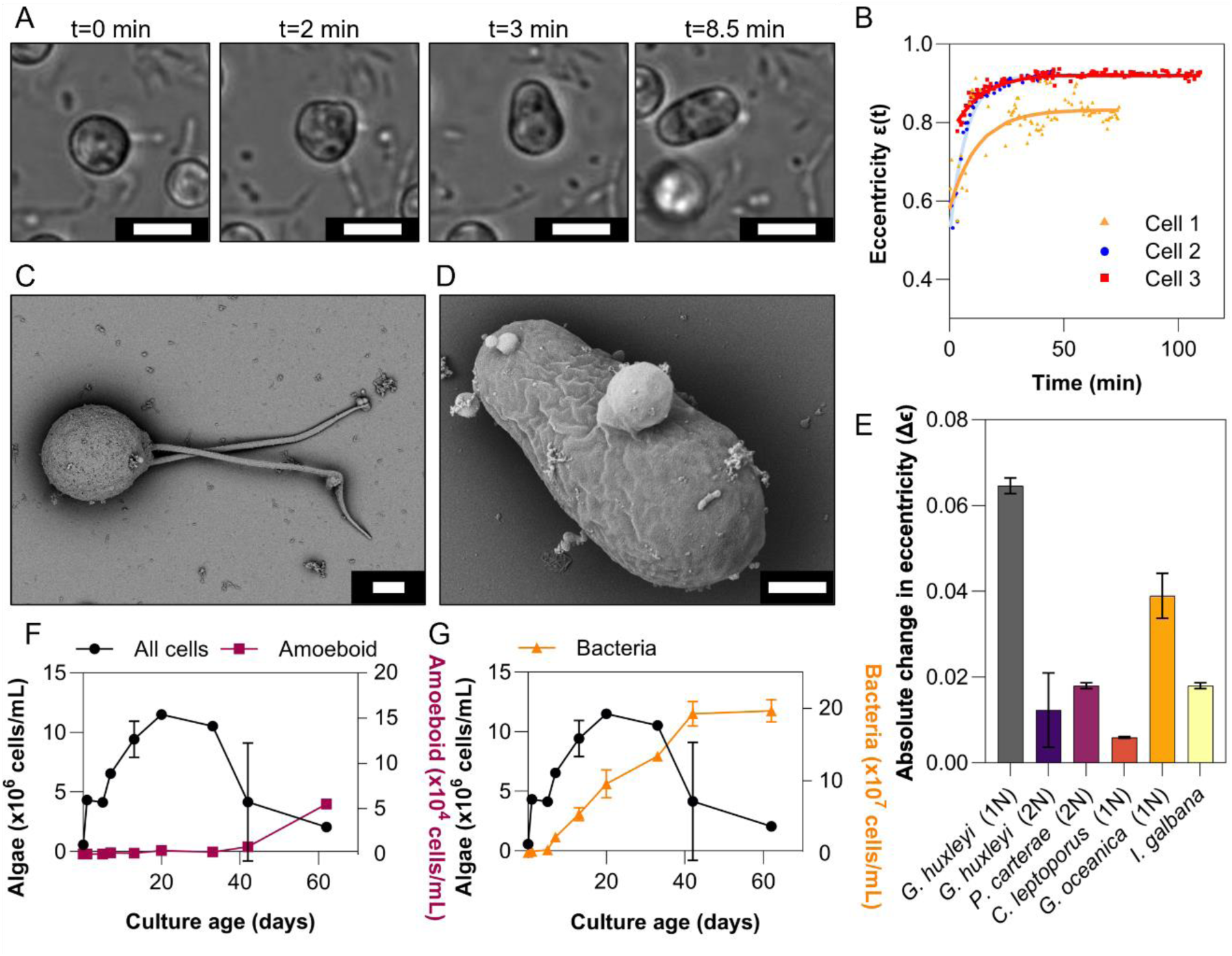
Metamorphosis haploid to amoeboid phase in coccolithophores. *(A)* Light microscopy time-lapse images of a haploid *G. huxleyi* cell in the transition to an ellipsoidal amoeboid cell (scale bar 5 µm). *(B)* Change in eccentricity (perfect sphere eccentricity ϵ = 0) over time for three cells (colored points) cell morphology transitions from broadly spherical to ellipsoidal with a characteristic timescale of 15 min (extracted from the exponential fits (lines)). *(C)* SEM image of a *G. huxleyi* haploid bi-flagellated cell (scale bar 1 µm). *(D)* SEM image of a *G. huxleyi* amoeboid cell (scale bar 1 µm). *(E)* Change in eccentricity after exposure to the bacteria, *Marinomonas* 5a1, and before exposure Δϵ = |ϵ(*treatment*) − ϵ(*control*)| for various haptophyte strains, with data plotted as the mean with standard error (SE) bars and n=3 biological replicates. *(F)* Cell counts as a function of time for an untreated xenic haploid *G. huxleyi* culture for all algal cells (black circles, left axis) and amoeboid cells (pink squares, right axis), with data plotted as the mean with SE error bars and n=3 biological replicates. *(G)* Cell counts as a function of time for xenic haploid *G. huxleyi* culture for all agal cells (black circles, left axis) and total bacterial cells (orange triangles, right axis), with data plotted as the mean with SE error bars and n=3 biological replicates.

We observed considerable amoeboid cell formation only in the haploid morphotypes of the two bloom-forming coccolithophores *G. huxleyi* and *G. oceanica*, and not for example in diploid *G. huxleyi* or in the closely related haptophyte *Isochrysis galbana* (Fig. 1E, SI Fig. S3). Monitoring *G. huxleyi* cultures via flow cytometry revealed that the amoeboid cells could be distinguished from the normal haploid population by a shift in forward scatter area (FSC-A) (SI Fig. S4). SYBR-Green DNA staining and flow cytometry showed that relative cellular DNA content of haploid and amoeboid populations is similar, suggesting that the transition from the haploid phase to the amoeboid phase conserved the ploidy of the original haploid phase (SI Fig. S4). The amoeboid phase becomes increasingly prominent (up to 3% of the original population) as cultures age into stationary and declining phase (Fig. 1F), consistent with previous anecdotal observations that amoeboid cells were observed in older cultures^16^. Furthermore, bacterial counts in xenic haploid *G. huxleyi* cultures, obtained by flow cytometry, show that the increase in amoeboid concentration coincides after the peak in bacterial cell number and begins to rise as the overall *G. huxleyi* culture collapses (Fig. 1G).

Metamorphosis into the amoeboid phase was accompanied by a considerable change in motility. Haploid cells swim in a meandering, looping trajectory with a rapidly decorrelating swimming direction, reflected in an average autocorrelation time of 2.8 ± 0.1 s and a mean speed of 26.3 ± 1.8 µm/s (Fig. 2). In contrast, amoeboid cells have an average autocorrelation time of 7.2 ± 0.9 s, thus changing direction approximately three times less often than haploid cells per unit time (Fig. 2B, Movie S2), with an average speed of 7.1 ± 0.1 μm/s (Fig. 2A), approximately a quarter of the average haploid swimming speed. Furthermore, the swimming speed distribution is narrower for the amoeboid cells in comparison to haploid cells, with standard deviations of 0.3 and 5.6 respectively. During motion the cell body can be clearly seen to rotate (Movie S2), which combined with the asymmetrical body shape gives the appearance of ‘wriggling’, as noted in previous records of amoeboid cells^36^. However, we were unable to resolve the flagella motion, so it remains unclear whether and how the flagella beat patterns are altered by this metamorphosis.

**Fig. 2.**
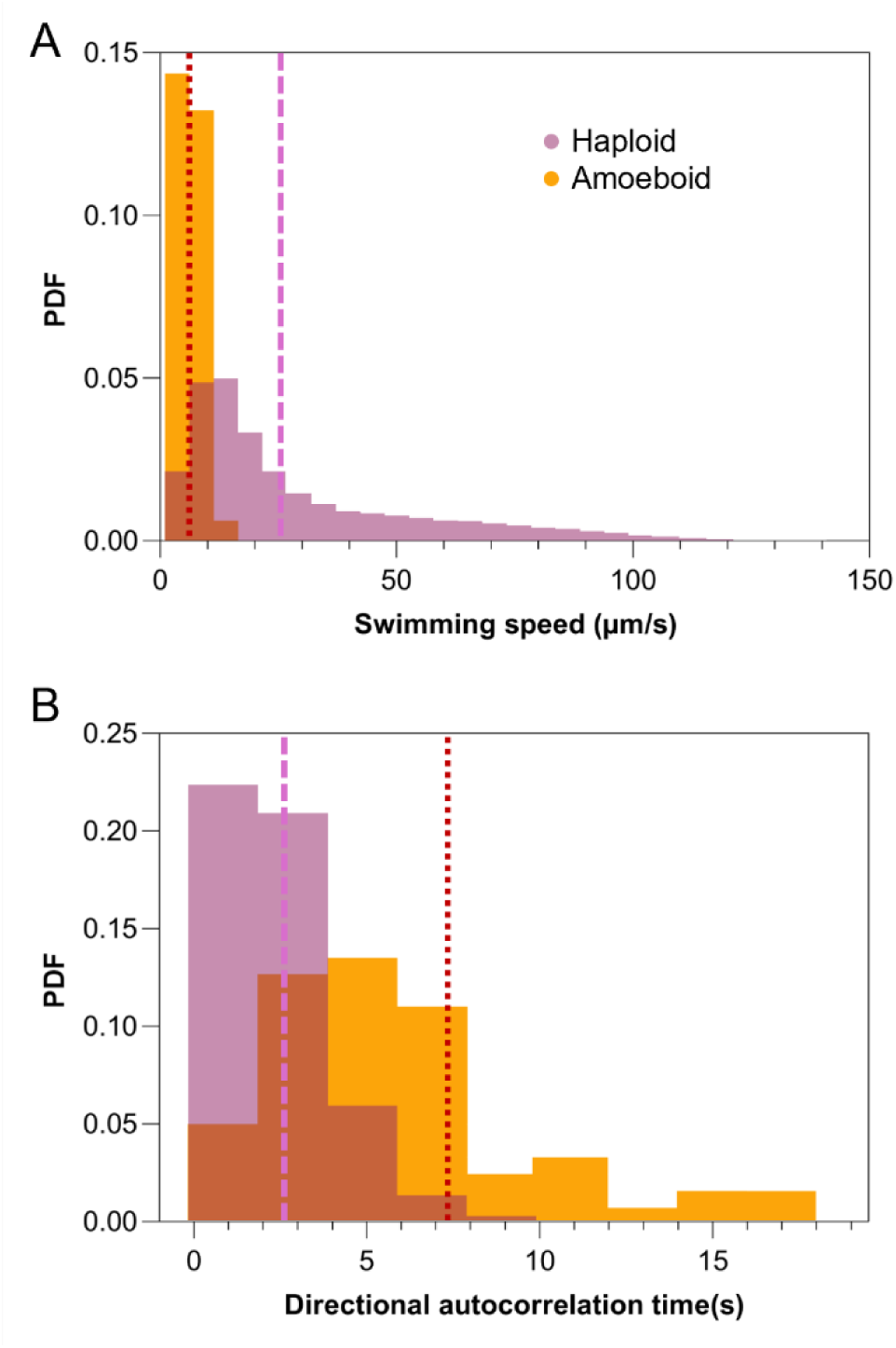
Motility of *G. huxleyi* haploid and amoeboid cells. *(A)* Probability density functions of swimming speed of *G. huxleyi* haploid cells (purple) and amoeboid cells (yellow) with speeds of 26.3 ± 1.8 μm/s and 7.1 ± 0.1 μm/s (mean ± standard error of the mean, SEM), respectively, with n = 8 biological replicates. (B). Velocity autocorrelation of the haploid and amoeboid phases, with an autocorrelation time of 2.8 ± 0.1 s and 7.2 ± 0.9 s (mean ± SEM) for the haploid and amoeboid respectively, with n=8 biological replicates. Average speeds and autocorrelation times indicated with dashed purple (haploid) and dotted orange (amoeboid) lines.

### Amoeboid cells maintain photophysiology and are not induced by typical environmental triggers

To assess the cell fitness of amoeboid cells, quantum yields were measured using amplitude modulation (PAM) microscopy. Measurements of the maximum quantum yield and relative electron transport rate (rETR) are commonly used as proxies for cell health and photosynthetic activity, respectively^41,42^. A comparison between haploid and amoeboid cells revealed no significant difference in either the maximum quantum yield (Fig. 3A) or rETR (Fig. 3B), indicating that cells maintain comparable photosynthetic fitness after transformation to the amoeboid phase^43,44^.

**Fig. 3.**
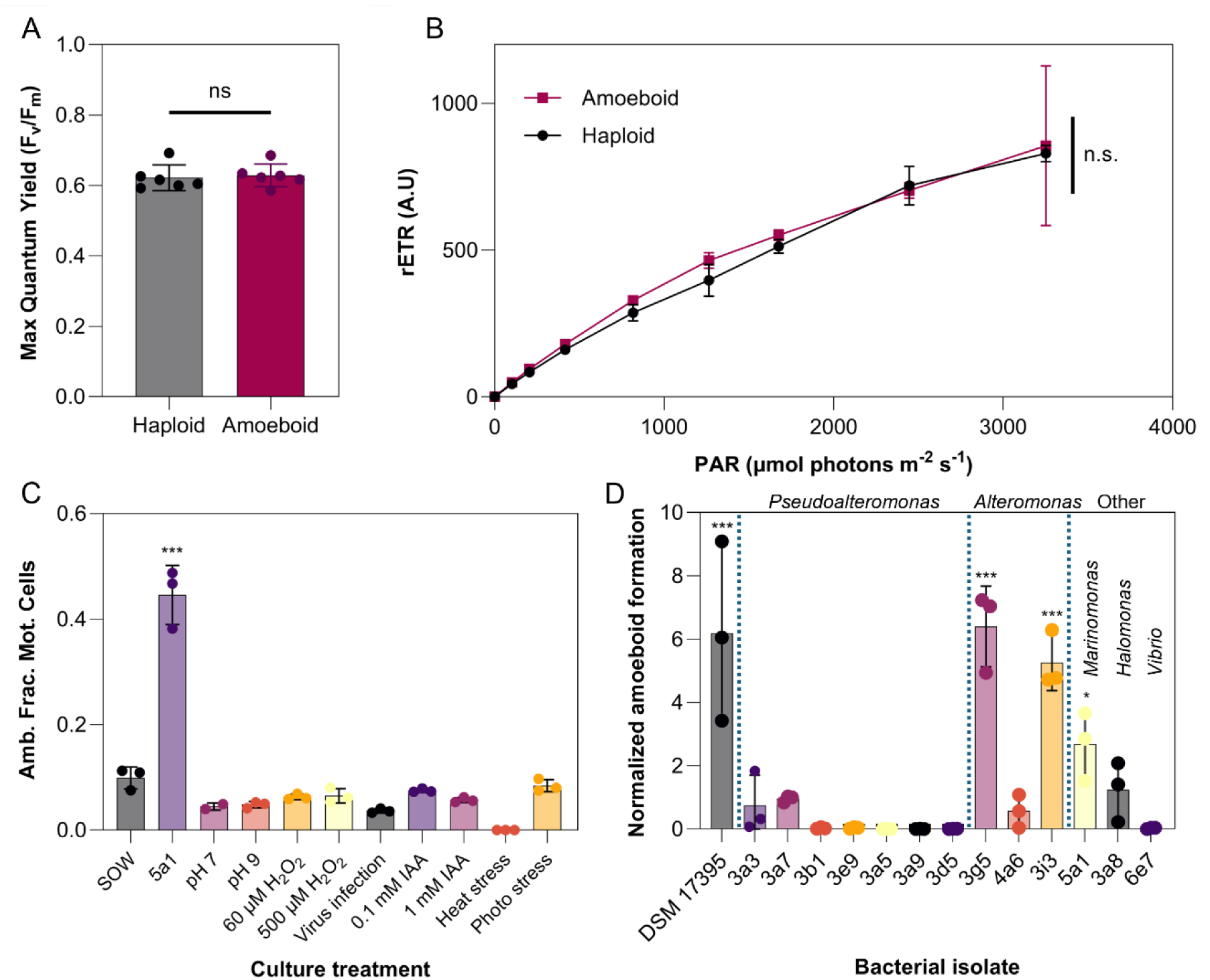
Amoeboid cells maintain photophysiology and metamorphosis is only triggered by exposure to environmentally-associated bacteria and not other environmental stressors. PAM measurements of haploid (gray) and amoeboid cells (purple, induced via exposure to bacterial strain *Marinomonas* 5a1), neither the *(A)* maximum quantum yield or *(B)* relative electron transport rate (rETR) differ between the amoeboid and haploid phases. Data plotted as the mean with SE error bars with n=6 biological replicates. A 2-tailed t-test showed no statistically significant differences (p-value < 0.05) between the two phases for both the maximum and effective quantum yields (across all PAR values). *(C)* The amoeboid fraction amongst motile cells (Amb. Frac. Mot. Cells) due to exposure of *G. huxleyi* haploid cells to different environmental stressors. Only exposure to bacteria (here the isolate *Marinomonas* 5a1) induced the formation of a significantly greater number of amoeboid cells in comparison with the synthetic ocean water (SOW) control (no stressors applied). Data shown as the mean with SE error bars and n=3 biological replicates. A one-sided ANOVA (p-value < 0.05) was carried out followed by a one-tailed post-hoc Tukey test applied for each treatment against the SOW (p-value < 0.05). *(D)* Relative amoeboid population, after exposure of haploid *G. huxleyi* cells to environmentally associated bacterial isolates^45^ from five bacterial genera: *Pseudoalteromonas*, *Alteromonas*, *Marinomonas*, *Halomonas* and *Vibrio*, and exposure to the *G. huxleyi* pathogen *Phaeobacter inhibens* (DSM 17395). Amoeboid induction is shown to be bacterial-strain dependent, with significant induction in response to 4 out of the 14 strains. Data normalized against the SOW treatment and plotted as the mean with SE error bars and n=3 biological replicates. A one-sided ANOVA (p-value < 0.05) was carried out followed by a one-tailed post-hoc Tukey test applied for each treatment against the SOW (p-value < 0.05).

We performed a systematic screening of common environmental stressors to determine the trigger of the amoeboid transformation. Xenic haploid cultures were exposed to different potential stressors (Methods), including rapid changes in pH, temperature increases, viral infection, oxidative stress, photo-induced stress, and indole-3-acetic acid (at either growth-enhancing (0.1 mM) or algicidal (1 mM) concentrations)^11^. Cultures were then maintained in pre-exposure growth conditions and monitored over 24 h (Fig. 3C). None of these stressors triggered a significant amoeboid response, indicating that the amoeboid phase transition is not a generalized stress response. Instead, a significant fraction of the population underwent metamorphosis into the amoeboid phase when a haploid culture was inoculated with high densities of an environmentally relevant bacterial strain, *Marinomonas* 5a1, previously isolated from a *G. huxleyi* bloom in a mesocosm experiment^45^ in Bergen (Norway) (Fig. 3C). This observation is consistent with our finding that amoeboid cells are induced within stationary phase xenic haploid cultures with a high density of bacteria (Fig. 1FG). The metamorphosis into the amoeboid phase also does not appear to be related to a change in predatory strategy: whilst the haploid cell contains acidic compartments for bacterial phagocytosis^46^, staining of both the diploid and amoeboid demonstrated the absence of such compartments (SI Fig. S5, Methods).

### Amoeboid formation is triggered by bacterial exposure resembling bloom collapse conditions

To examine the specificity of the impact of bacteria on the transformation from haploid to amoeboid phase, haploid *G. huxleyi* cultures were independently inoculated with 13 bacterial strains isolated from the same mesocosm experiment in Bergen^45^. Amoeboid cells were characterized through light microscopy and particle tracking (Methods) and identified through their differences in motility and morphology (Fig. 1CD, Fig. 2AB). Aside from the above identified *Marinomonas* 5a1 strain (Fig. 3C), two *Alteromonas* strains (3g5, 3i3) also triggered considerable amoeboid induction in *G. huxleyi* (Fig. 3D). No response was observed to any of the seven isolated *Pseudoalteromonas* strains. Significant amoeboid induction was further observed with the known *G. huxleyi* pathogen *Phaeobacter inhibens* (DSM 17395, found in the microbiome of *G. huxleyi* isolated from the Galician coast)^11,37,47,48^. These observations demonstrate a specificity in the metamorphosis trigger, with only particular coccolithophore-associated bacterial strains inducing amoeboid formation.

To investigate the nature of the algal–bacteria interaction responsible for triggering the transformation, *Marinomonas* 5a1 was used to prepare three treatments. Overnight cultures of 5a1 were washed and incubated in (i) unsupplemented Synthetic Ocean Water (SOW) medium, (ii) a stationary phase haploid *G. huxleyi* culture, and (iii) filtered medium from a stationary phase haploid culture, respectively. After 24 h of incubation, the three cultures were filtered to remove all cells and the filtrates were then supplied to three stationary-phase haploid cultures of *G. huxleyi*. The first and third treatments triggered amoeboid formation (Fig. 4A), demonstrating that the presence of bacterial exudates alone is sufficient to induce transformation, without the need for physical contact between the bacteria and algae. In contrast, filtrates from the second treatment did not induce any significant amoeboid formation, suggesting that potential chemical triggers were either degraded or consumed by the algae during the incubation stage.

**Fig. 4.**
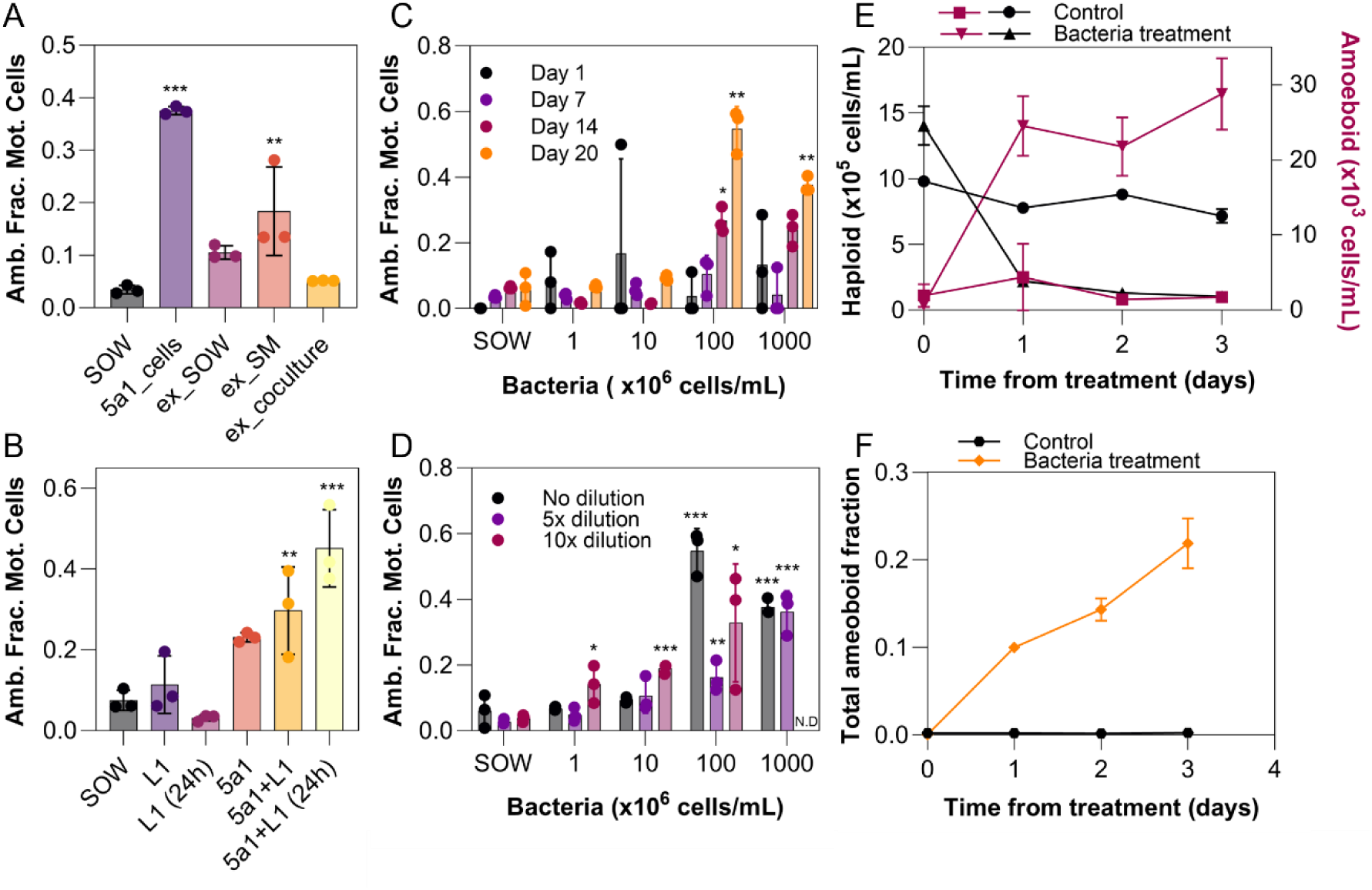
Algal and bacterial conditions required for bacterial-induced amoeboid formation in *G. huxleyi*. *(A)* The fraction of amoeboid cells among motile cells (Amb. Frac. Mot. Cells) upon exposing haploid *G. huxleyi* cultures to exudates generated from three different treatments of *Marinomonas* 5a1: synthetic ocean water (SOW), spent media of *G. huxleyi* and *G. huxleyi* culture. (*B)* The fraction of amoeboid cells among motile cells under different nutrient conditions: control condition (SOW), addition of inorganic L1 nutrients (phosphorus, nitrate, silicate, trace metals, and vitamins) added either immediately prior to incubation with 5a1 or 24 h prior to the start of the experiment. *(C)* The fraction of amoeboid cells among motile cells at four time points of algal culture (day 1, black, pre-exponential phase; day 7, purple, exponential phase; day 14, pink, stationary phase; day 20, orange, declining phase) for different supplemented bacterial concentrations. Significant amoeboid induction occurred only in stationary and older *G. huxleyi* cultures with bacterial concentrations exceeding 10^8^ cells/mL. *(D)* The fraction of amoeboid cells among motile cells in *G. huxleyi* haploid cultures (starting cell concentration of 10^6^ cells/mL) induced upon varying algal:bacterial cell ratios. Significant amoeboid induction occurred at higher bacterial concentrations, but is also seen at lower bacterial concentrations when the algal culture is itself diluted. (*A-D*) All data shown as the mean with SE error bars and n = 3 biological replicates. A one-sided ANOVA (p-value < 0.05) was carried out followed by a one-tailed post-hoc Tukey test applied for each treatment against the SOW (p-value < 0.05). (*E* and *F*) Flow cytometry counts (*E*) of haploid and amoeboid *G. huxleyi* cells in a control condition (black circles, pink squares) and cultures co-incubated with 5a1 (black triangles, pink inverted triangles). (*F*) Resulting fraction of amoeboid cells from a control culture (black circles) and a culture inoculated with *Marinomonas* 5a1 (orange diamonds). Data shown as the mean with SE error bars and n = 3 biological replicates.

Further experiments indicate that the transformation is not a starvation response. Inoculating stationary phase haploid cultures with both *Marinomonas* 5a1 and a nutrient supplement (containing organic nutrients, phosphate, nitrate, silicate, trace metals, and vitamins, so as to supplement the medium back to starting concentrations of the L1 algae growth medium; Methods) still resulted in amoeboid formation (Fig. 4B). This indicates that metamorphosis is not induced by the consumption of the initially supplied inorganic nutrients by either bacteria or algae.

Induction of the amoeboid phase primarily occurs in conditions typical of those following a bloom collapse. Amoeboids were originally significantly detected with high densities of both algae and bacteria (10^6^ cells/mL and 10^9^ cells/mL, respectively). To test for a dependence of amoeboid metamorphosis on algal cell age, four different ages (1, 7, 14, and 20 days old) of haploid *G. huxleyi* cultures were inoculated with bacterial cultures of different concentrations, from 10^6^ to 10^9^ cells/mL (Fig. 4C). Significant amoeboid induction was observed in cultures older than 14 days (i.e., stationary and older, SI Fig. S6) in the two highest starting bacterial concentrations (10^8^ and 10^9^ cells/mL). In a further experiment, a 20-day old haploid culture with starting density of 10^6^ cells/mL was divided into triplicate cultures and diluted by either 10x, 5x, or no dilution, then inoculated with different bacterial concentrations (Fig. 4D). Amoeboid formation in these diluted *G. huxleyi* cultures was observed for initial bacterial concentrations of 10^6^ cells/mL and higher. Taken together, these results identify two important aspects of the metamorphosis. Firstly, amoeboid formation is not induced in young cultures regardless of starting bacterial densities. Secondly, the formation of amoeboid cells at algal concentrations of 10^5^ cells/mL and bacterial concentrations of 10^6^ cell/mL, matching concentrations reported in naturally occurring *G. huxleyi* blooms^23,49^, indicates that this third life phase may be ecologically relevant in the ocean in late-stage algal blooms.

Tracking both the amoeboid and haploid phases in *G. huxleyi* over time with flow cytometry (Methods) showed that exposure to *Marinomonas* 5a1 resulted in a small sub-population, consisting of approximately 3% of the original haploid population, transitioning to amoeboid cells within three days (Fig. 4E). This sub-population persisted under the bacteria-rich conditions that induced it, whereas the cell count of the initial haploid population simultaneously decreased by 93% within three days (Fig. 4E). This resulted in in 21% of the final algal population being represented by amoeboid cells (Fig. 4F). Taken together with our observations that the metamorphosis occurs under conditions closely resembling the demise of an algal bloom, these results suggest that the amoeboid morphology could be a survival mechanism of *G. huxleyi* against the rise of bacteria in the final stages of an algal bloom.

## Discussion

In this study, we have characterized another life phase in bloom-forming coccolithophores. While this life phase was previously reported anecdotally^16,37,38^, both a detailed description and the identification of the conditions required to trigger its occurrence had been lacking. Our experiments show that upon exposure to certain bacterial strains, the haploid phase of *G. huxleyi* undergoes a rapid morphological transformation, over a timescale of 15 min, during which the cell elongates from a nearly spherical to a 3:1 ellipsoid (Fig. 1AB) and the average swimming speed reduces by 73% (Fig. 2A). In this new amoeboid phase, PAM measurements show that cells retain the same photophysiology as the haploid cells (Fig. 3AB), indicating that amoeboids maintain comparable photosynthetic fitness to the original haploids. We find that a suite of environmental stressors, including inorganic nutrient starvation (Fig. 4B), viral infection, oxidative stress, pH changes and photostress all fail to induce this metamorphosis (Figs. 4B, 3C), while several species of bacteria do. This transformation does not require any physical contact between algae and bacteria (Fig. 4A). In these conditions the amoeboid population persists after the haploid population crash, reaching up to 21% of the total cell numbers (Fig. 4D-F). Given that the Marinomonas isolate and some of the bacterial species tested are known to have algicidal effects on coccolithophores^11,47,48^, we hypothesize that the persistent morphotype may confer resistance to bacteria.

The conditions that trigger the transition from haploid to amoeboid phase are consummate with those of the harsh conditions generated by algal blooms^23,50–53^. Firstly, induction was only observed in bloom-forming coccolithophore species (*G. huxleyi* and *G. oceanica*) and not in other common coccolithophores, including the closely related but non-bloom-forming haptophyte *Isochrysis galbana* (Fig. 1E). Secondly, this metamorphosis occurs predominantly in older stationary/declining haploid cultures in the presence of high densities of certain bacterial strains, including a well-known *G. huxleyi* pathogen and strains isolated from a mesocosm in which a coccolithophore bloom had been induced^45^ (Fig. 2D). The population fraction of amoeboids in unsupplemented xenic cultures also correlates with size of the bacterial population (Fig. 1FG), consistent with prior observations of amoeboids in stationary phase cultures^16^.

The reason why this metamorphosis occurs remains unclear. The rise of amoeboids as a prominent population fraction during late-bloom-like conditions, together with the simultaneous demise of the haploid cells from which amoeboids derive, suggests that this metamorphosis could be a late-stage survival response. On the other hand, the rapid transition could also be a farming mechanism^11,54^, that is a transformation elicited by the bacteria to harvest algal exudates more efficiently or for a long time after the haploid phase has succumbed to the bacterial-induced conditions.

Regardless of the root cause, the transition into a third life phase expands the underlying mechanisms driving algal blooms and their collapse, particularly as the transition into the amoeboid phase is not the first instance of coccolithophores adapting to rapidly-changing environmental conditions. The general progression of bloom dynamics in coccolithophores has been well documented^23,49,55^: it starts with an increase in the diploid population and is followed by a rise in viral particles that lyse the majority of the diploid cells, leading to the formation of a viral-resistant sub-population of haploid cells^33–35^. The large release of organic matter through viral lysis supports a bacterial bloom^56,57^. Our results point to a natural continuation of this sequence, where the rapid bacterial growth triggers the metamorphosis of the haploid cells into the amoeboid phase. There are thus clear similarities between the viral-induced transition from diploid to haploid, and the bacterial-induced transition from haploid to amoeboid.

A natural question prompted by our work is how the coccolithophore life cycle proceeds after the amoeboid phase. Whilst we have not observed a transition from the amoeboid to either the haploid or diploid phases, this would close the new expanded life cycle of *G. huxleyi.* If such a transition exists, it likely requires specific environmental triggers that have so far proved elusive under laboratory conditions. For example, by isolating individual amoeboid cells away from the bacterial population or completely replacing the media, we have not observed a reversal of the metamorphosis. Further work will also be required to determine whether the transition to the amoeboid phase is directly triggered by the bacteria or is an active defense response against the bacteria, which signaling factors drive it, and what biomechanical mechanism underpins the shape shift. We have shown that amoeboids, unlike haploids, persist in bacteria-rich conditions (Fig. 4E), and that there are surface changes associated with the metamorphosis (SI Fig. S2), which could possibly impair bacterial recognition of this new phase^58–60^. Another possible explanation is the exudate composition of amoeboid cells could be shifted compared to haploid exudates, thus avoiding detection from bacteria, something that would require a dedicated metabolomic investigation.

We expect the amoeboid phase to be more prevalent than the lack of reports to date would imply. It would be intriguing and further cement the ecological relevance of amoeboids to search for signs of this life phase in existing environmental field data such as Imaging FlowCytobot data (IFCB)^61^. The robustness of the transformation makes it appear likely that amoeboid cells have been overlooked due the drastic morphological differences compared to the traditionally studied diploid and haploid cells. Given the rise of algal bloom events linked with ongoing rising ocean temperatures^62,63^, we anticipate these results will promote future investigations into the inter-species interactions shaping algal bloom dynamics and further our understanding of these key microbial players on the global scale.

## Materials and Methods

### Algal strains and growth conditions

All algae strains were grown in L1 supplemented synthetic ocean water (SOW, pH = 8)^64^. Haploid and diploid *G. huxleyi* strains RCC1217 and CCMP3266 (also known as RCC1216) respectively, *Gephyrocapsa oceanica* haploid strain RCC1315, *Isochrysis galbana* strain RCC1353 and *Pleurochrysis carterae* diploid strain RCC6317 were cultured at 18 °C on a 14:10 h diurnal cycle at a light intensity of 100 µmol m^-2^ s^-1^. *Calcidisucus leptoporus* haploid strain RCC1131 was cultured at 18 °C on a 14:10 h diurnal cycle at a light intensity of 60 µmol m^-2^ s^-1^. *Coccolithus braarudii* diploid strain RCC1200 was cultured at 14 °C on a 12:12 h diurnal cycle at a light intensity of 30 µmol m^-2^ s^-1^.

### Bacterial growth and amoeboid induction conditions and exudate preparation

Bacterial strains were previously isolated from an induced mesocosm *G. huxleyi* bloom in Bergen^45^ and preserved in 20% glycerol solution at −80 ⁰C. Bacteria were inoculated in Marine Broth media (Difco, CAS 10043-52-4) from frozen stocks and grown overnight in a shaking incubator at 30 ⁰C with constant agitation (200 revolutions per minutes, RPM). Cultures were collected at optical density (OD) 1.70–1.80 and were washed and resuspended three times (4000 relative centrifugal force, RCF, for 5 min) with SOW. Algal cultures were then inoculated with the washed bacterial cultures at a cell ratio of 100:1 (unless otherwise stated), and incubated at the relevant temperature, diurnal light cycle, and light intensity for each of the algae strains tested (specified above, see *Algal strains and growth conditions*) for 24 h to induce amoeboid formation.

Exudates were prepared by first growing the isolated strain 5a1 as described above, then inoculating three different treatments with a concentration of 10^9^ cells/mL (measured with flow cytometry, see *Flow cytometry and staining*): (i) fresh SOW medium, (ii) filtered (0.2 µm filter, Filtropur) spent medium from stationary *G. huxleyi* haploid culture, and (iii) a stationary *G. huxleyi* haploid culture. All treatments were incubated at 18 ⁰C on a 14:10 h light:dark cycle at 100 µmol m^-2^ s^-1^ for 24 h. After incubation, all bacterial and algal cells were removed via filtration (0.2 µm filter, Filtropur), and the resultant filtrate inoculated into replicate algal cultures (see *Algal strains and growth conditions*) with 1:10 *G. huxleyi* culture: exudate volume ratio (Fig. 4A).

### Preparation of algal stressors

*G. huxleyi* lytic virus EhV-201^65^ were generously provided by Prof. Assaf Vardi and maintained in exponential phase axenic diploid *G. huxleyi* (CCMP3266) cultures. *G. huxleyi* haploid cultures were infected with EhV-201 at multiplicity of infection (MOI) of one^66,67^. High and low pH conditions (pH values of 9 and 7, respectively) were created by the addition of NaOH and HCl to haploid cultures, respectively. *G. huxleyi* haploid cultures were thermally stressed by rapidly raising the culture temperature from 18 ⁰C to 50 ⁰C for 10 min. Photostress was imposed by subjecting the cells to epi-illumination at 587 nm light supplied from a Sola Light Engine Lumencor at full power (∼4W) equipped to an inverted microscope (Nikon Ti-E; 10× 0.3 NA objective and mCherry filter cube). Oxidative stress was induced by adding hydrogen peroxide (60 µM) and indole-3-acetic acid (IAA, Merck, CAS 87-51-4) was prepared in 50% ethanol-water solutions and diluted to final concentrations of 0.1 and 1 mM IAA in the haploid *G. huxleyi* culture with a final ethanol concentration of 1%^48^.

### Microscopy and single-cell tracking

Imaging chambers were prepared by separating one standard glass microscopy slide (Avantor VMR) and a standard cover glass slide (Number 1.5, Epredia) with a 1 mm thick gasket fabricated by punching a polydimethylsiloxane (PDMS; Dow Corning SYLGARD 184) sheet with an 8 mm diameter biopsy punch. Samples were gently pipetted into the imaging chambers prior to sealing with the cover glass slide. Once loaded, imaging chambers were incubated at 18 ⁰C for 24 h on a 14:10 h light:dark cycle at 100 µmol m^-2^ s^-1^. Imaging of the samples was performed using phase-contrast microscopy (Nikon Ti-E; 10× 0.3 NA objective). Videos for cell tracking and shape analysis were recorded at 25 frames per second using a CMOS camera (Orca-Flash 4.0, Hamamatsu).

Cell tracking was conducted using TrackPy (v.0.6.1)^68^, described previously for tracking bacterial movement^45^. The background of each image was removed by computing the absolute difference of each pixel value with the median intensity value per pixel over the full length of the video (30 s, 600 frames). Particle detection was performed on the absolute intensity difference with a particle size of 17 pixels and a minimum distance of 51 pixels between particles. For the assembly of trajectories a maximum displacement of 17 pixels and maximum disappearance of 2 frames was allowed. Resulting cell velocities were calculated with a five-frame moving-average. Trajectories shorter than 10 frames were discarded. For the autocorrelation analysis, the cellular positions were processed with a second-order Savitzky-Golay filter with a time window of 5 frames and the change in angle was computed from the filtered positional data. The autocorrelation function was then computed using the Python Numpy function correlate, on the angle time series of each trajectory. The correlation function was then fit to a single exponential function to obtain the autocorrelation time of the directional persistence. Cells were classified as amoeboid provided the average velocity < *v* > ≤ 8 μ*m*. *s*^−1^ based off the speed distributions (Fig. 2A) and an eccentricity ϵ ≥ 0.8.

### Scanning electron microscopy and energy-dispersive X-ray spectroscopy

200 µL liquid cultures of haploid and amoeboid cells were fixed with 1% (w/v) glutaraldehyde and deposited on hydrophilized 0.01% poly-L-lysine coated silicon wafers. After 10 min incubation, the wafers were sequentially immersed for 5 min each in 2.5% glutaraldehyde solution (SOW, 27 practical salinity units, PSU), 1% osmium tetroxide, and SOW. Wafers were then passed through an ethanol drying series by sequential immersion in 0%, 30%, 50%, 70%, 90%, and 100% ethanol for 2 min at each stage, and finally three times in water-free ethanol. Wafers were dried using a critical point dryer with a cell monolayer protocol (CPD 931 Tousimis, ETH Zurich Microscope Facility ScopeM) and were fixed to aluminum SEM stubs using silver paint. Samples were degassed for 24 h then sputter coated with 4 nm platinum-palladium (CCU-010 Metal Sputter Coater Safematic, ETH Zurich Microscope Facility ScopeM).

Scanning electron microscopy (SEM) imaging and energy dispersive X-ray spectroscopy (EDX) were performed using an extreme high resolution (XHS) TFS Magellan 400 microscope (ETH Zurich Microscope Facility ScopeM) with 50 pA current and an accelerating voltage of 5 kV for imaging and 15 kV for EDX. Scattered electrons were collected and imaged with both an immersion backscattered electron CBS detector and secondary electron TLD detector. EDX was performed on the amoeboid cells and heavily calcified *Coccolithus braarudii*, rather than diploid *G. huxleyi* due to the gradual decalcification of laboratory-maintained *G. huxleyi* cultures^11^.

### Microbial adhesion to hydrocarbons assay

*G. huxleyi* adhesion to hydrocarbons assay was performed based on a previously described protocol^69^. *G. huxley*i cells of RCC1217 haploid were grown to stationary phase and amoeboid metamorphosis was induced as described previously (see Bacterial growth and amoeboid induction conditions and exudate preparation). Amoeboid and haploid cultures were filtered with a 1 µm cell filter (pluriStrainer, pluriSelect), then washed and resuspended with Phosphate urea magnesium sulphate (PUM) buffer^69^ with cell concentrations adjusted to 30×10^3^ algae cells/mL. 150 µL of n-hexadecane was then added to 5 mL glass vials containing 1.5 mL cell suspension and vortexed for 2 minutes. These samples were allowed to settle for 15 min and then the number of cells in the aqueous hexadecane free layer were measured using the flow cytometry protocol with APC-A (see *Flow cytometry and staining*). These cell counts were then compared to cell counts from control blanks without hexadecane for both cultures. The percentage hydrophobicity was then calculated as follows: Hydrophobicity (%) = (A_0_-A_1_)/ A_0_ x 100, where A_0,1_ are the algal cell counts in the aqueous layers of hexadecane-free blank and hexadecane-treated samples respectively.

### Pulse amplitude modulation microscopy (PAM)

An amoeboid culture was prepared by inoculating a stationary phase *G. huxleyi* haploid culture with a washed and resuspended *Marinomonas* 5a1 culture (see Bacterial growth conditions and exudate preparation) and incubated at 18 ⁰C on a 14:10 light:dark cycle at 100 µmol m^-2^ s^-1^ for 24 h prior to imaging. A 30 µL sample of cell suspension was confined within a silicone gasket (Grace Bio Labs, 0.5 mm thickness) separating two coverslips (Epredia, Number 1.5) coated with 0.1% (w/v) poly-L-lysine solution (Sigma Aldrich, CAS 25988-63-00) to immobilize the cells. The cells were then dark adapted for 15 min prior to being loaded onto a chlorophyll-fluorescence imaging microscope^70^ (Florescence Kinetic Microscope; Photon Systems Instruments), with bright-field and fluorescence images captured with a TOMI 2 camera (Photo Systems Instruments). Bright-field images were used to identify cell positions for extraction of fluorescence parameters from the chlorophyll fluorescence images using a custom python script. The distributions of the maximum and effective quantum yields showed a single population peak, confirming that the majority of the cells were of one type and not that two separate populations in the amoeboid treatment averaged to the same value (*SI Appendix Fig. S6*). The maximum quantum yield of photosynthetic energy conversion 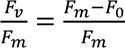 and the effective quantum yield of photosynthetic energy conversion 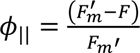 where measured using the saturation pulses method^41,71^, where *F*_*v*_, *F*_*m*_, and *F*_0_ are the variable, maximal and minimal fluorescence of dark adapted cells, respectively, *F* the fluorescence emission of dark-adapted cells, and *F*_*m*_′ the maximal fluorescence of light-adapted cells. Rapid light curves were constructed by measuring effective quantum yields at increasing levels of photosynthetically active radiation (PAR). The maximum quantum yield of photosynthetic energy conversion in PSII provides a relative measure of the maximal efficiency at which light drives PSII photochemistry. The relative transport of electrons (rETR) driven by photosystem II (PSII) was calculated as *rETR = PAR X ϕ_||_*, which estimates the relative rate of realized linear electron flow through PSII at a given PAR intensity.

### Flow cytometry and staining

Algal cells were counted using a flow cytometer (Beckman Coulter, CytoFLEX S) equipped with a 488 nm laser and forward scatter (FSC) and red (allophycocyanin APC-A) fluorescence was recorded for each sample for a standardized time per sample of 40 s. Bacterial and algae cultures for determining ploidy were stained and incubated with SYBR Green (5 µM final concentration, Sigma Aldrich CAS 163795-75-3) for 10 min in the dark and counted using a flow cytometer recording FSC and green (fluorescein, FITC) fluorescence for a standardized time of 60 s per sample. Amoeboid cultures were first filtered with a 1 µm cell filter (pluriStrainer, pluriSelect) and resuspended in filtered media prior to counting to prevent machine clogging.

### Cells acidic compartment staining

*G. huxleyi* cells (RCC1217 haploid and CCMP3266 diploid) were grown to stationary phase and amoeboid metamorphosis was induced overnight in RCC1217 as described above (see *Bacterial growth and amoeboid induction conditions and exudate preparation*). 1 mL samples were collected from haploid, diploid and amoeboid phases and stained with Lysotracker^TM^ Deep Green DND-26 (ThermoFischer Scientific, L7526) to a final concentration of 50 nM^72^. After staining, cells were imaged immediately using a fluorescent microscope (Nikon Ti-E; 30× X NA objective, Sola Light Engine Lumencor) and a FITC filter cube (Chroma 49002).

## Acknowledgments

We thank Assaf Vardi, Assaf Gal, Jean-Baptiste Raina, Martin Ackermann, Jonasz Słomka, Zachary Landry, Frédéric de Schaetzen, Dieter Baumgartner and Isobel Short for helpful discussions, Russell Naisbit for scientific editing and feedback, and Ela Burmeister for assistance in experimental work. We thank Samuel Charlton for assisting in testing surface modifications. We thank Assaf Vardi for gifting of the EhV-201 viral strain. The authors gratefully acknowledge Stephan Handschin, Miriam Lucas, and Anne Greet Bittermann of ScopeM at ETH Zurich for their support & assistance in this work.

## Supplementary Information

**Supplementary Figure 1.**
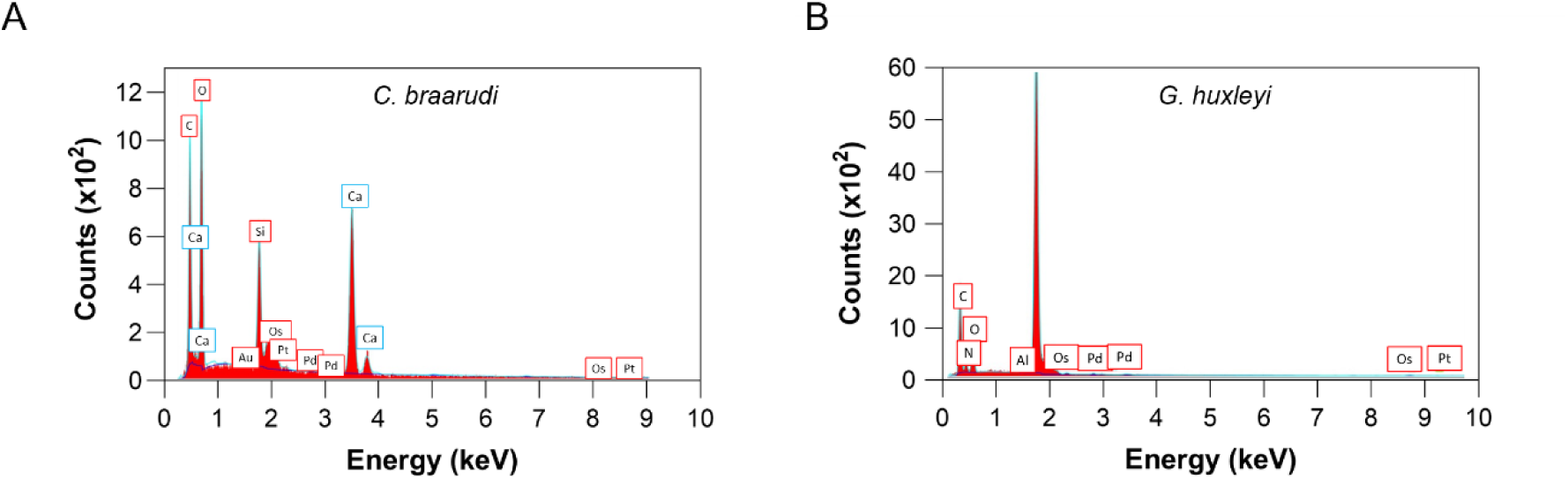
Energy-dispersive X-ray (EDX) spectra of diploid *Coccolithus braarudii* and amoeboid *Geopharycapsa huxleyi*. (A) Characteristic peaks of calcium carbonate, indicating cell calcification in *C. braarudii* but not in (B) *G. huxleyi*.

**Supplementary Figure 2.**
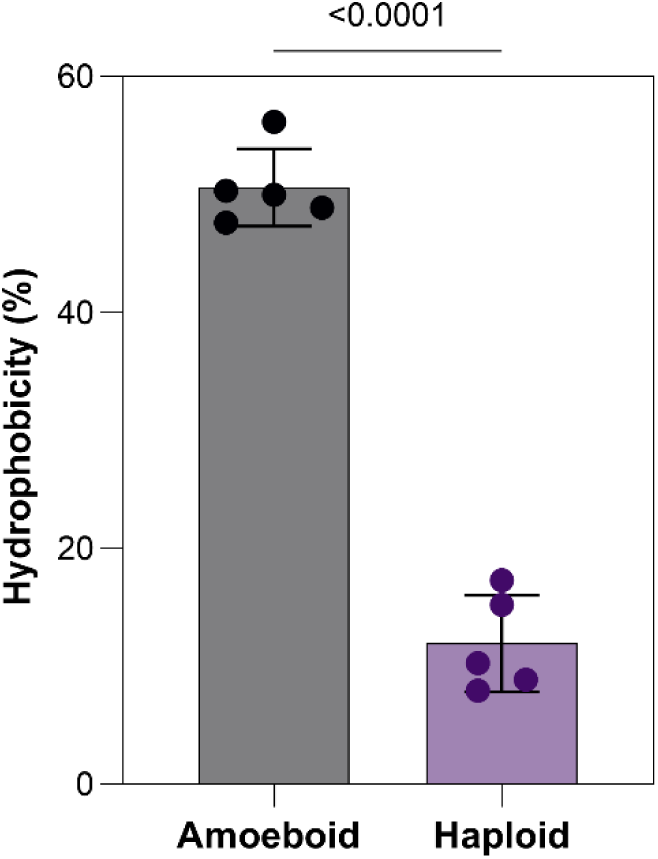
Comparison of surface hydrophobicity between *G. huxleyi* haploid and amoeboid phases. A microbial adhesion to hydrocarbon (MATH) assay was performed on haploid and amoeboid *G. huxleyi* revealing an increase of 39 ± 2% hydrophobicity in the amoeboid phase. Data shown as the mean with n=5 biological replicates with SE error bars. A 2-tailed t-test was applied for the hydrocarbon-free treatment against the blank hydrocarbon-free treatment (p < 0.05).

**Supplementary Figure 3.**
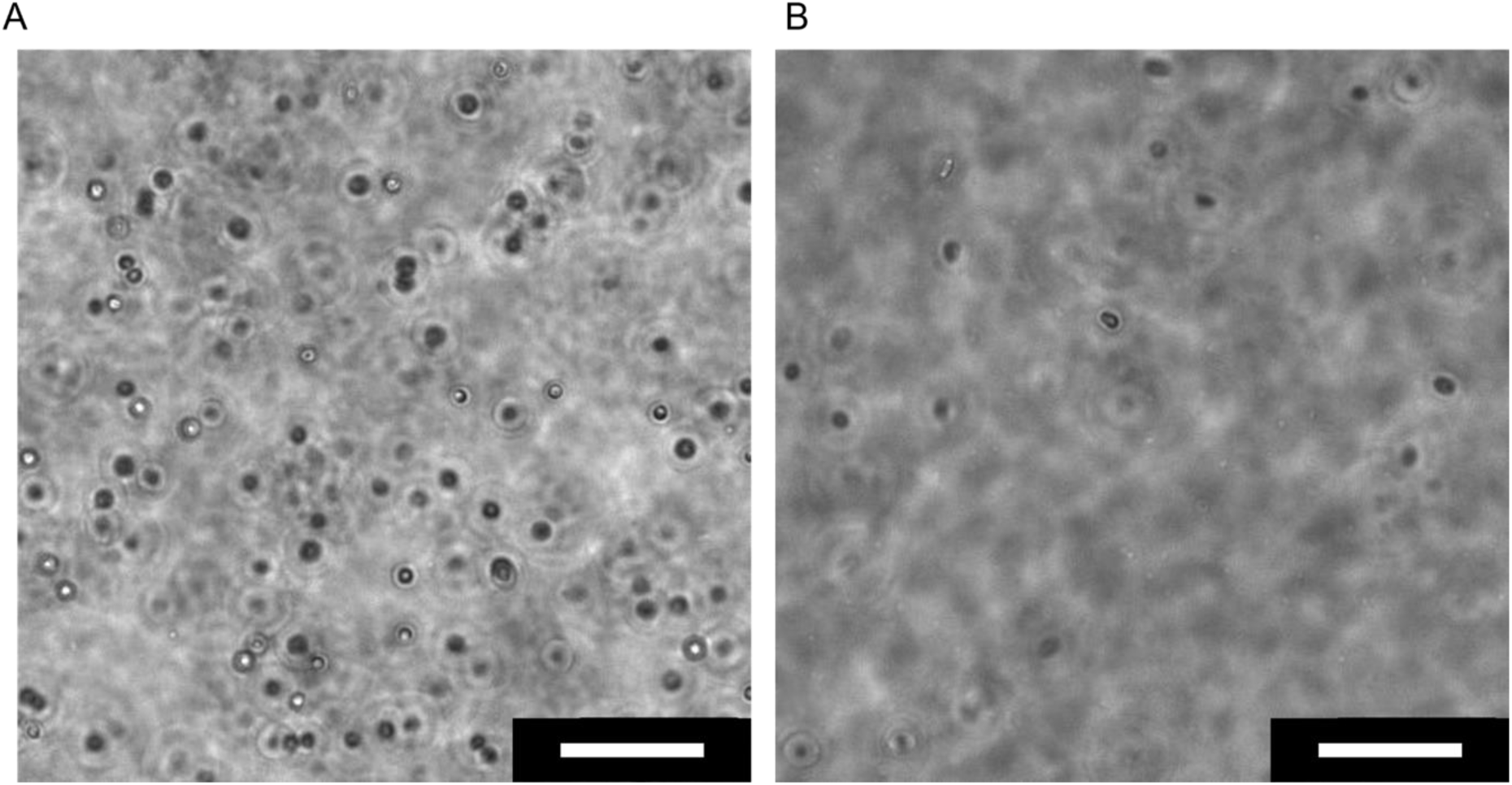
Light microscopy images of (*A*) haploid and (*B*) amoeboid *Geopharycapsa oceanica.* (A) In control conditions haploid *G. oceanica* cells show a spherical morphology. (B) In treatment conditions where haploid *G. oceanica* cultures were exposed to *G. huxleyi*-associated bacteria, the algal cells underwent a morphological transformation from spherical to an elongated cell morphology. Scale bars represent 50 μm.

**Supplementary Figure 4.**
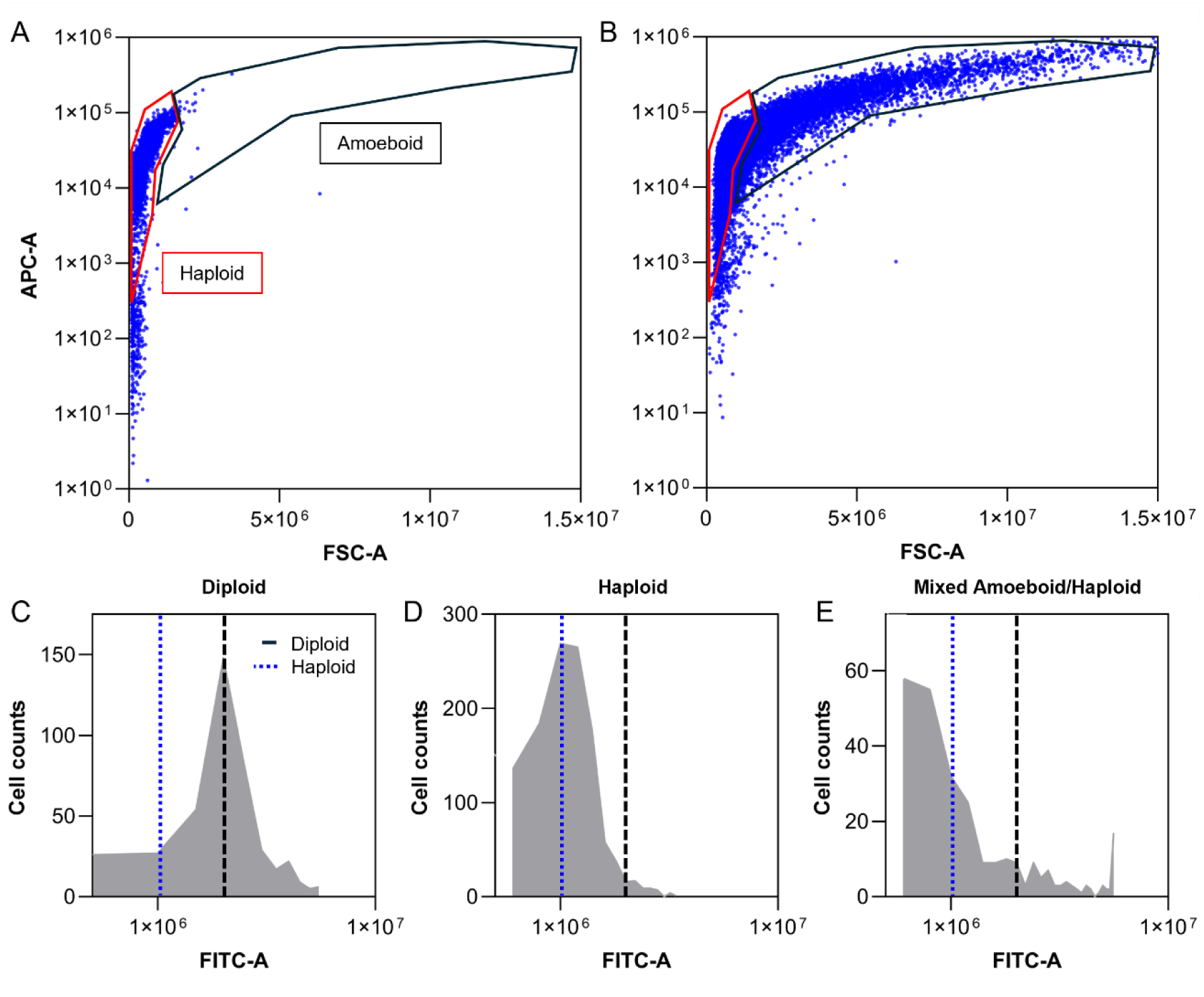
Flow cytometry gating for the identification of *G. huxleyi* amoeboid cells and FITC-A distributions for ploidy determination. Autofluorescence chlorophyll APC-A fluorescent signal for *G. huxleyi* (A) haploid and (B) mixed haploid and amoeboid cultures. Ploidy was determined from the FITC-A fluorescence distribution signal obtained from SYBER green staining of *G. huxleyi* cultures, identifying diploid/haploid characteristic FITC-A intensities (black dashed line and dotted blue lines respectively) for the following *G. huxleyi* cultures: (C) diploid, (D) haploid, and (E) mixed amoeboid and haploid.

**Supplementary Figure 5.**
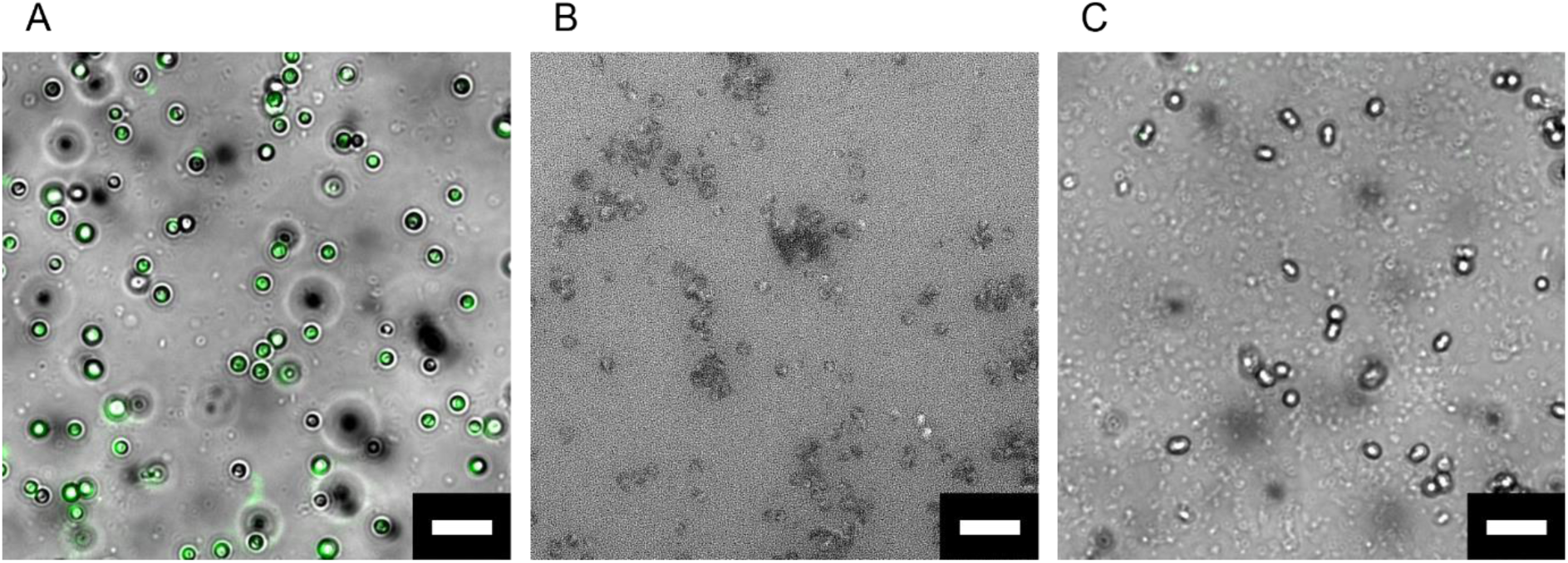
Fluorescence microscopy images of *G. huxleyi* cells stained for the acid compartment using LysoTracker^TM^ Deep Green DND 26. Staining of the acidic compartment reveals signal in the haploid phase (*A*) but not in the diploid *(B)* or the amoeboid *(C)* phases, consistent with reports of phagocytosis by haploids^46^. The lack of staining in diploid *G. huxleyi* confirmed that the acidic compartments involved in calcification were not stained through this method and therefore the staining observed in the haploid (*A*) was not due to these acidic compartments^73,74^. Scale bars represent 20 μm.

**Supplementary Figure 6.**
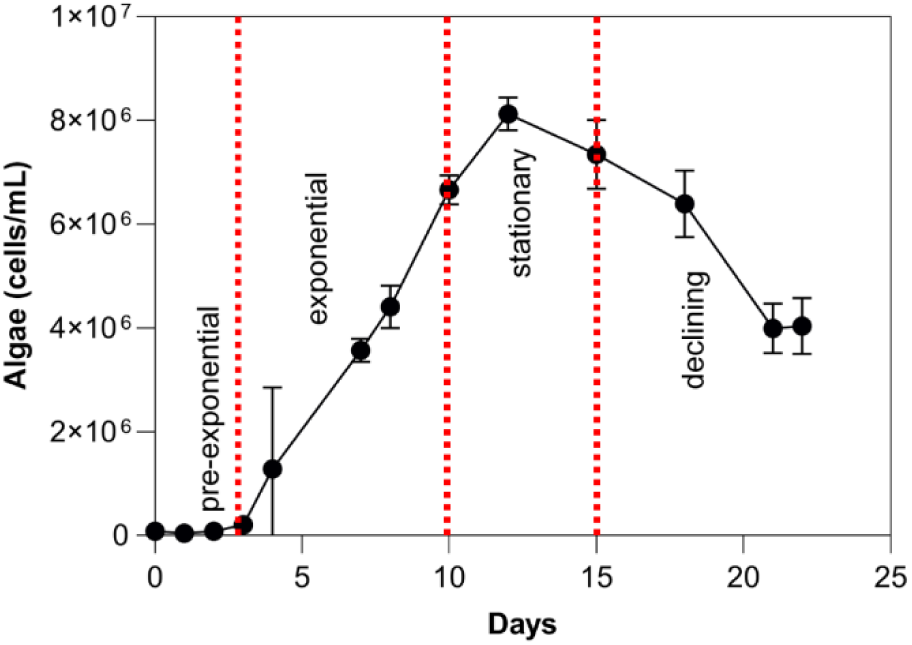
Growth curve of *Geopharycapsa huxleyi* RCC1217. Cell counts of haploid *G. huxleyi* cultures were measured using flow cytometry (see Methods) to identify the growth phases. Data shown as the mean with SE error bars and n=3 biological replicates.

**Supplementary Figure 7.**
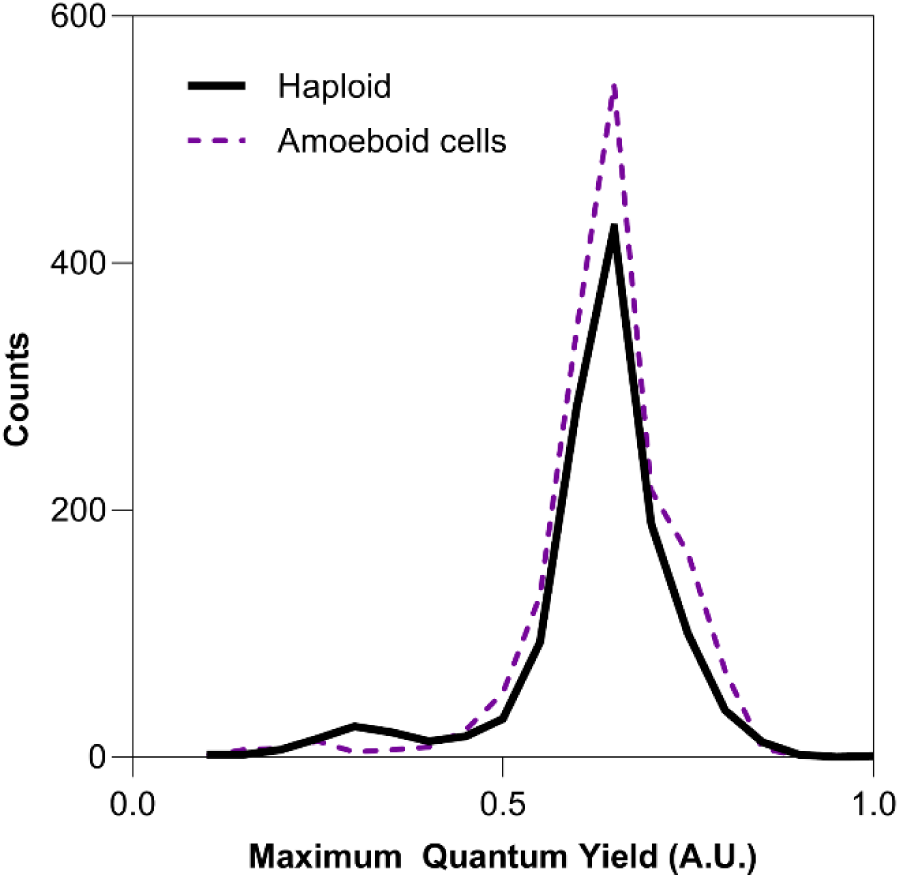
Maximum quantum yield distribution curves for control and bacteria-treated *G. huxleyi* cultures. Maximum quantum yields were obtained using Pulse Amplitude Modulation (PAM) microscopy with analysis areas selected using light microscopy to determine amoeboid-enriched areas. The absence of a significant second peak in either of the conditions confirms that there was a single dominant population of cells.

**Supplementary Video S1. Transformation of haploid *Geopharycapsa huxleyi* into the amoeboid morphotype.** Timelapse microscopy video (two frames per min) showing transformation of a spherical haploid *G. huxleyi* cell into an elongated “amoeboid” cell type over a period of approximately 16 min.

**Supplementary Video S2. Swimming of amoeboid *Geopharycapsa huxleyi.*** Light microscopy video (20 fps) of *G. huxleyi* culture treated with bacteria 24 h prior to imaging, illustrating the clear cell body rotation of the amoeboid cell.

